# The effects of hitchhiking and repulsion of deleterious alleles on linkage disequilibrium during a selective sweep

**DOI:** 10.64898/2026.10.01.756034

**Authors:** Maya Weissman, Kirk E. Lohmueller, Nandita R. Garud

## Abstract

Deleterious mutations are abundant in nature. Consequently, when an adaptive mutation arises, it will likely land on a haplotype with several deleterious mutations. While deleterious mutations may hitchhike with the adaptive variant, deleterious mutations are less likely to be found on the same haplotype with each other due to repulsion (i.e. Hill Robertson interference). Both hitchhiking and repulsion shape patterns of linkage disequilibrium (LD), but their simultaneous impact on LD have not been fully understood, leaving a gap in accurate inference of selection from genomic data. Here, we investigate patterns of LD during a sweep using simulations across a wide range of evolutionary parameters. On average, LD is elevated between pairs of intermediate frequency nonsynonymous variants relative to synonymous variants. This occurs because nonsynonymous variants, which are presumably deleterious, are more likely to rise to high frequency via hitchhiking rather than drift and thus will be predominantly found on the sweeping haplotype. However, we find that this elevation does not occur across all distances: when nonsynonymous variants are either too close to each other or too far from the adaptive locus, repulsion between deleterious variants results in lower LD among nonsynonymous variants. This results in a characteristic “criss-crossing” of LD. We find that criss-crossing is a feature of empirical data from *Clostridioides difficile*, a pathogen that has undergone several selective sweeps. However, some sweeps show elevation of nonsynonymous LD over synonymous LD across all distances. These sweeps have been previously identified to bear complex adaptations, and moreover, harbor elevated nucleotide diversity. Consistent with this, we demonstrate that lack of criss-crossing can arise when adaptation proceeds via epistatic, multi-locus architecture or introgression.

## 1 Introduction

Adaptation is a process whereby beneficial mutations rise to high frequency within a population. As an adaptive allele sweeps to high frequency, nearby linked variants will also “hitchhike” to high frequency. ^1^ Consequently, hitchhiking is known to lead to regions of locally elevated linkage disequilibrium (LD), a signal which has been widely leveraged to identify selective sweeps from genomic data. ^2,3^

However, the extent of LD in the vicinity of a selective sweep in part depends on the selective effect of linked variants (e.g. deleterious or neutral). Recently it has been found that LD among pairs of intermediate frequency (i.e. minor allele frequency *>* 0.2) nonsynonymous variants 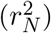 is on average elevated relative to pairs of intermediate frequency synonymous variants 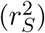 at selective sweeps. ^4^ Since nonsynonymous variants are more likely than synonymous variants to be deleterious, they are less likely to reach high frequency by drift alone, and instead may reach high frequency due to hitchhiking. ^5,6^ Consequently, initially rare nonsynonymous variants that hitchhike during a sweep will be largely private to the adaptive haplotype, generating strong LD between pairs of nonsynonymous variants. By contrast neutral synonymous variants can rise to high frequency via both drift and hitchhiking, and so can be found on adaptive and non-adaptive haplotypes, generating weaker LD.

Hitchhiking is not the only force that can result in differential LD patterns among nonsynonymous vs. synonymous variants. Previous work has shown that deleterious variants experience repulsion due to Hill Robertson interference^7,8^ and negative synergistic epistasis. ^9–11^ Hill Robertson interference specifically occurs when two non-interacting deleterious variants co-occur at very short distances between each other, as the probability that the variants can be broken apart by recombination is low, and thus purifying selection against haplotypes bearing both mutations increases. ^12–14^ While the previous works have demonstrated depressed mean LD between rare deleterious nonsynonymous variants relative to synonymous variants due to Hill Robertson interference, ^7,8^ the combined effect of repulsion with hitchhiking on LD has not yet been explored.

Here, we investigate patterns of 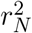 vs. 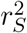 during a sweep in simulations spanning a wide range of evolutionary parameters. We find that while 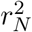 is on average greater than 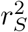 during selective sweeps, 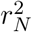 is less than 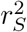 at short distances between variants. We show that this “criss-crossing” of LD arises from the competing forces of hitchhiking, which elevates 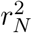 greater than 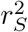 at moderate to large distances, and repulsion, which depresses 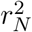 less than 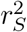 at short distances. We also demonstrate that hitchhiking is able to mask the effects of repulsion on LD when deleterious pairs are in very close proximity to the sweeping beneficial variant. This criss-crossing pattern is common across a wide range of parameters, as well as empirically in *Clostridium difficile*. However, a subset of selective sweeps in *C. difficile* do not exhibit criss-crossing, and instead show 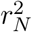 greater than 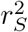 across all distances. These sweeps have been previously shown to be complex adaptations ^15,16^ and moreover show spikes in nucleotide diversity. Using simulations, we find that the lack of criss-crossing can arise under simple models of multi-locus, epistatic interactions or adaptive introgression. Taken together, our results characterize how deleterious variation and modes of adaptation jointly shape patterns of LD during a selective sweep.

## 2 Materials and methods

### 2.1 Simulations

We performed forward population genetics simulations in SLiM 4. ^17^ We simulated a panmictic population of *N*_*e*_ = 10^4^ individuals and a chromosome of length 10^4^ bp. Unless specified, mutation rate was *µ* = 10^*−*6^. Nonsynonymous mutations were subject to a selection coefficient of *s*_*del*_ as specified in the text and synonymous mutations were neutral. The dominance coefficient, *h*, of all mutations was held constant at 0.5, unless specified. The ratio of synonymous to non-synonymous mutations was 1 : 2.3. Unless specified, recombination occurred with probability *r* = 10^*−*6^ per base pair. Following a burn-in of 10 *\* N*_*e*_ generations, beneficial nonsynonymous mutations, with selection coefficient *s*_*ben*_, were permitted to evolve at a single focal site with mutation rate proportional to *θ*_*ben*_ = *Ne * µ*_*ben*_ = 0.01, unless specified (e.g. in the case of a soft sweep). Simulations terminated when the focal beneficial mutation reached a frequency of 0.5. At the conclusion of each simulation, a random sample of 100 individuals was drawn to calculate LD.

#### 2.1.1 Demographic contraction models

We also modeled demographic contractions in the absence of a selective sweep. Simulations again began with a burn-in with population size *N*_*e*_ = 10^4^ for 10 *\* N*_*e*_ generations, at which point an instantaneous demographic contraction occurred. In the “short, sharp contraction” model, the population size was reduced to 1*/*10 *\* N*_*e*_ and simulations continued for an additional 20 generations. In the “long, shallow contraction” model, the population size was reduced to 1*/*2 *\* N*_*e*_ and the simulations continued for an additional 5000 generations.

#### 2.1.2 Model of adaptive introgression and epistasis

To model epistatic adaptation and adaptive introgression, we simulated an adaptive fragment with multiple loci (*n*_*loci*_) conferring the beneficial selective effect. Regardless of whether we modeled epistasis or introgression, when all *n*_*loci*_ mutations are present on the same haplotype, the adaptive fragment has a total selection coefficient of *s*_*together*_ = 0.1. In the epistasis model, interacting nonsynonymous variants were each deleterious on their own (*s*_*alone*_ = *s*_*del*_ = *−*0.001) but became beneficial (*s*_*together*_*/n*_*loci*_ = 0.01) when all *n*_*loci*_ variants were combined on the same haplotype. In the adaptive introgression introgression model, only the single, central nonsynonymous variant is beneficial (*s*_*alone,ben*_ = *s*_*together*_ *−* (*n*_*loci*_ *−* 1) *\* s*_*alone,del*_), while the remaining *n*_*loci*_*−*1 nonsynonymous variants are deleterious hitchhikers with *s*_*del*_ = *−*0.001.

In both our models of epistasis and introgression, following the neutral burn-in, 5 individuals were randomly selected to simultaneously receive all *n*_*loci*_ nonsynonymous variants in the adaptive fragment. Individuals were selected to receive the full adaptive fragment to reduce computation time, as the waiting time to acquire *n*_*loci*_ = 10 deleterious mutations is long. The distance between each nonsynonymous variant and the next in the fragment was drawn from a Poisson distribution with *λ* = 10 bp. The mutation rate for all nonsynonymous variants in the fragment was *µ*_*ben*_ = 0, such that the only way for mutations in the adaptive fragment to become unlinked was through recombination. Simulations terminated when the center mutation of the adaptive fragment, but not necessarily the entire fragment, reached a frequency of 0.5.

### 2.2 Summarizing LD

We measured LD using the standard *r*^2^ statistic:

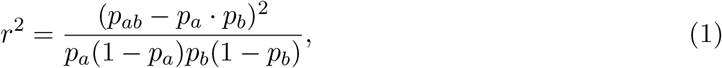

where *p*_*a*_ is the frequency of haplotypes bearing the variant at locus 1, *p*_*b*_ is the frequency of haplotypes bearing the variant at locus 2, and *p*_*ab*_ is the frequency of haplotypes bearing both variants. To enrich for positively selected alleles and hitchhikers, we focused on LD among variants that are intermediate frequency in the broader population (minor allele frequency 0.2). We computed LD between pairs of nonsynonymous variants 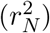 and synonymous variants 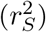.

To identify when pairs are positively correlated (i.e. linked) or negatively correlated (i.e. in repulsion), we calculate the signed correlation coefficient as:

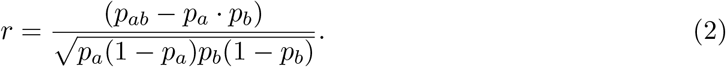

The signed correlation coefficient ranges from −1 to 1, where *r <* 0 denotes pairs that are negatively correlated, or are less likely to appear together than predicted by random chance, and *r >* 0 indicates pairs that are positively correlated, or are more likely to co-occur than predicted by random chance.

### 2.3 Analysis of empirical *Clostridium difficile* genomes

We analyzed 134 *C. difficile* isolate genomes from the Unified Human Gastrointenstinal Genome (UHGG) project (alias MGYG-HGUT-02369).^18^ We downloaded alignments of isolate genomes, as well as accompanying genomic data files used to annotate sites and genes, from MGnify. ^19^ We used custom scripts to annotate each bi-allelic SNP as synonymous/nonsynonymous. ^4^

#### 2.3.1 Detecting sweeps using iLDS

To detect selective sweeps, we recapitulated the integrated Linkage Disequilibrium Score (iLDS) scan of the *Clostridium difficile* genome from Wolff and Garud ^4^ (fig. 4). In brief, iLDS is calculated in sliding windows across the genome. Within each window, iLDS quantifies the difference in area under the LD decay curves for 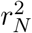 and 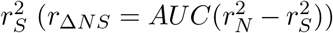. Additionally, iLDS scans for windows with locally elevated LD, *rL*^2^, compared to the genome-wide average, *rG*^2^ (*r*_Δ*LG*_ = *AUC*(*rL*^2^ *− rG*^2^)). Both 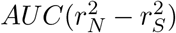 and *AUC*(*rL*^2^ *− rG*^2^) are normalized by the mean and standard deviation of these statistics calculated in sliding windows genome-wide 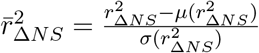, and then squared, giving the final statistic of:

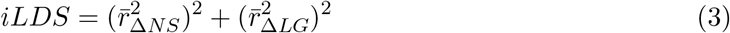

Windows were classified as significant by iLDS if both 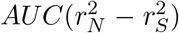 and *AUC*(*rL*^2^ *− rG*^2^) are significantly greater than 0 and the total iLDS score exceeds a critical threshold (see Wolff and Garud ^4^). Following Wolff and Garud ^4^, for *C. difficile* iLDS was applied in windows corresponding to approximately 10^4.5^ base pairs, as this was the distance at which LD decays among intermediate frequency nonsynonymous sites for *C. difficile*.

#### 2.3.2 Calculating average nucleotide diversity

Average nucleotide diversity was calculated in sliding windows across the *C. difficile* genome. First, for each locus, nucleotide diversity was calculated as:

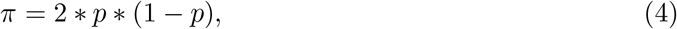

where *p* is the minor allele frequency at that locus. Individuals with missing data at a particular locus were omitted from the *π* calculation for that site. Next, the mean *π* per 1000 basepairs was calculated across sliding windows of 1000 bp and a step size of 500 bp between windows.

Genome-wide mean *π* was calculated by excluding all windows within a sweep.

#### 2.3.3 Calculating mean 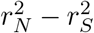 for empirical sweeps in *C. difficile*

We calculated both mean 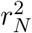 and 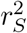 for all intermediate frequency (*MAF >* 0.2) variants (*MAF >* 0.2) within 10^4.5^ bp of the central locus of each of the six sweeps identified by iLDS. 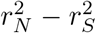was binned at distances of 30, 100, 500, 1000, 10000, 32000 such that there were at least 5 pairs per bin.

## 3 Results

### 3.1 Selective sweeps generate a ‘criss-crossing’ pattern of synonymous vs nonsynonymous LD

To assess patterns of LD during a selective sweep, we performed forward population genetics simulations in SLiM 4. ^17^ To enrich for selected alleles and their hitchhikers, we analyzed LD between pairs of intermediate frequency (*MAF >* 0.2) nonsynonymous variants 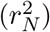 and pairs of intermediate frequency synonymous variants 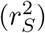 (fig. S1) using the standard *r*^2^ statistic (Methods).

First, to gain intuition about the expected signatures of LD in the absence of a sweep, we simulated panmictic populations of size *Ne* = 10^4^ with varying strengths of selection on the deleterious hitchhikers (*s*_*del*_). When all variants are neutral (*s*_*del*_ = 0), there is no difference in 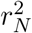 and 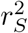 (fig. 1B). This can more readily be seen by plotting 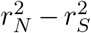, which is zero across all distances (fig. 1D, fig. S2).

**Figure 1.**
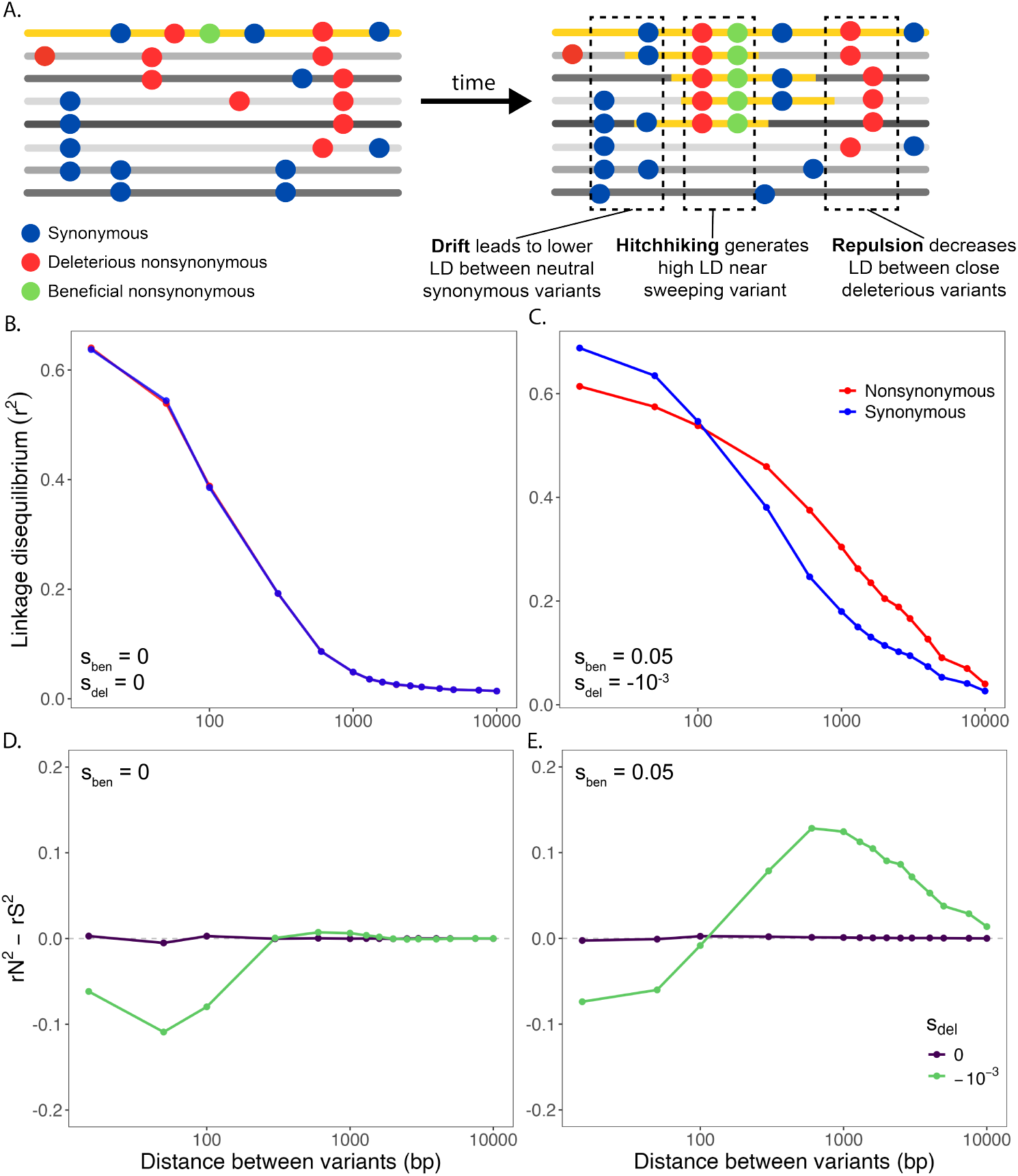
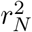 vs. 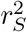 under neutrality compared to following a selective sweep. A. When a beneficial nonsynonymous variant (green) arises and sweeps to high frequency, linked synonymous (blue) and deleterious nonsynonymous (red) variants on the same haplotype can “hitchhike” to high frequency. Hitchhiking can lead to high LD between nonsynonymous variants near the sweeping beneficial locus. However, further from the selective sweep, Hill Robertson interference between deleterious variants can lead to lower 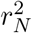 at short distances. B-E. Mean 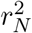 and 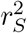 calculated from 1,000 replicate simulations of (B, D) neutrality (*s*_*ben*_ = 0) and (C, E) selection (*s*_*ben*_ = 0.05). Here, *Ne* = 10^4^, *r* = 10^*−*6^, *µ* = 10^*−*6^, *s*_*del*_ = *−*10^*−*3^, and *θ*_*ben*_ = 0.01. In C, criss-crossing of 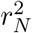 and 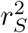 can be observed. (D,E) Criss-crossing of LD can more readily be seen by plotting the difference in 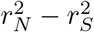. For additional evolutionary scenarios, see Figure 2 and Supplemental Figure 2.

However, when selection against deleterious variants can be felt (|*s*_*del*_| *>* 1*/Ne*), we find that at short distances between variants 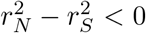 (fig. 1D, fig. S2). For example in fig. 1D, when *s*_*del*_ = *−*10^*−*3^, 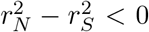 when pairs are *<* 300 bp apart. As the distance between variants increases, 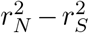 approaches, but does not exceed, 0. This occurs because the probability that two deleterious variants close together are unlikely to be broken apart by recombination, and thus selection against haplotypes bearing both mutations increases (fig. 1A), resulting in Hill Robertson interference. ^12–14^ Others have shown that Hill Robsertson interference will generate on average 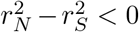 at short distances among rare deleterious variants. ^7,8^ Here, we find that this phenomenon can manifest even among intermediate frequency deleterious variants that may have risen to high frequency by chance despite being deleterious.

Next we asked whether hitchhiking of deleterious variants during a selective sweep could shift the mean of 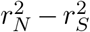 enough to offset the negative values produced by repulsion. Previously, we showed that deleterious variants can hitchhike to high frequency along with a beneficial variant, resulting in not only an increase in overall LD but also 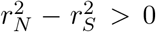. ^4^ We recapitulate this result: on average, 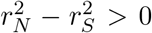 0 when 1*/Ne <* |*s*_*del*_| *<* |*s*_*ben*_| (fig. 1C, E, fig. S2). However, at short (e.g. *<* 100 bp) distances between variants, we observe 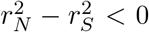, indicating that the effects of hitchhiking on LD are insufficient to overcome the effects of repulsion. A sweep therefore produces a characteristic “criss-crossing” of LD: 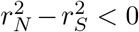 at short distances, where repulsion between deleterious variants has larger impact on mean LD, crossing to 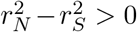 at intermediate distances, where hitchhiking has a larger impact (fig. 1C, E). As shown in the next section and in Supplemental Figures 2, 3, and 4, this criss-crossing signature is robust across a wide range of evolutionary parameters. We note that this criss-crossing effect is only observable when using finer binning at short distances than the distance at which 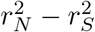 crosses 0 (fig. S6).

Finally, we tested whether these patterns of 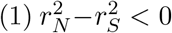 at short distances and 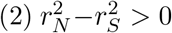 at intermediate distances can alternatively arise from other, non-selective forces. Demographic contractions do not generate any appreciable increase of 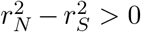 at intermediate distances, though repulsion still produces 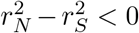 at short distance (fig. S5). Because allele frequency can impact LD, ^20,21^ we confirmed that 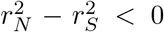 at short distances and 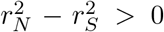 at intermediate distances was not sensitive to subtle differences in the distribution of allele frequencies between synonymous and nonsynonymous variants (fig. S7).

### 3.2 Criss-crossing occurs across a range of evolutionary parameters

Criss-crossing requires 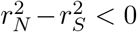 at short distances and 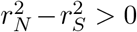 at intermediate distances. In this section, we find that the criss-crossing signature is a robust feature of selective sweeps across a broad range of evolutionary parameters with a few exceptions. We further show that the distance at which the crossover occurs and the magnitude of the effects on either side of it both depend on the underlying parameters.

First, criss-crossing manifests when *s*_*ben*_ is sufficiently strong (e.g. *>* 10^*−*2^) and |*s*_*ben*_| *>* |*s*_*del*_|(fig. 2A, fig. S2). As the strength of *s*_*ben*_ increases, the effect of hitchhiking increases, resulting in an increase in the maximum height of the 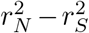 curve and a decrease in the distance at which the 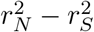 line crosses 0. Notably, even in the strongest regime tested (*s*_*ben*_ = 0.5), repulsion is still sufficient to produce a negative region at short distances.

Second, the strength of *s*_*del*_ impacts both the maximum height of 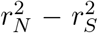 at intermediate distances and the depth of the decrease in 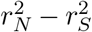 at short distances (fig. 2B). We observe the biggest shifts away from 0 with intermediate strength of selection. Moreover, *s*_*del*_ impacts where criss-crossing occurs, with the criss-crossing distance increasing with the strength of *s*_*del*_.

**Figure 2.**
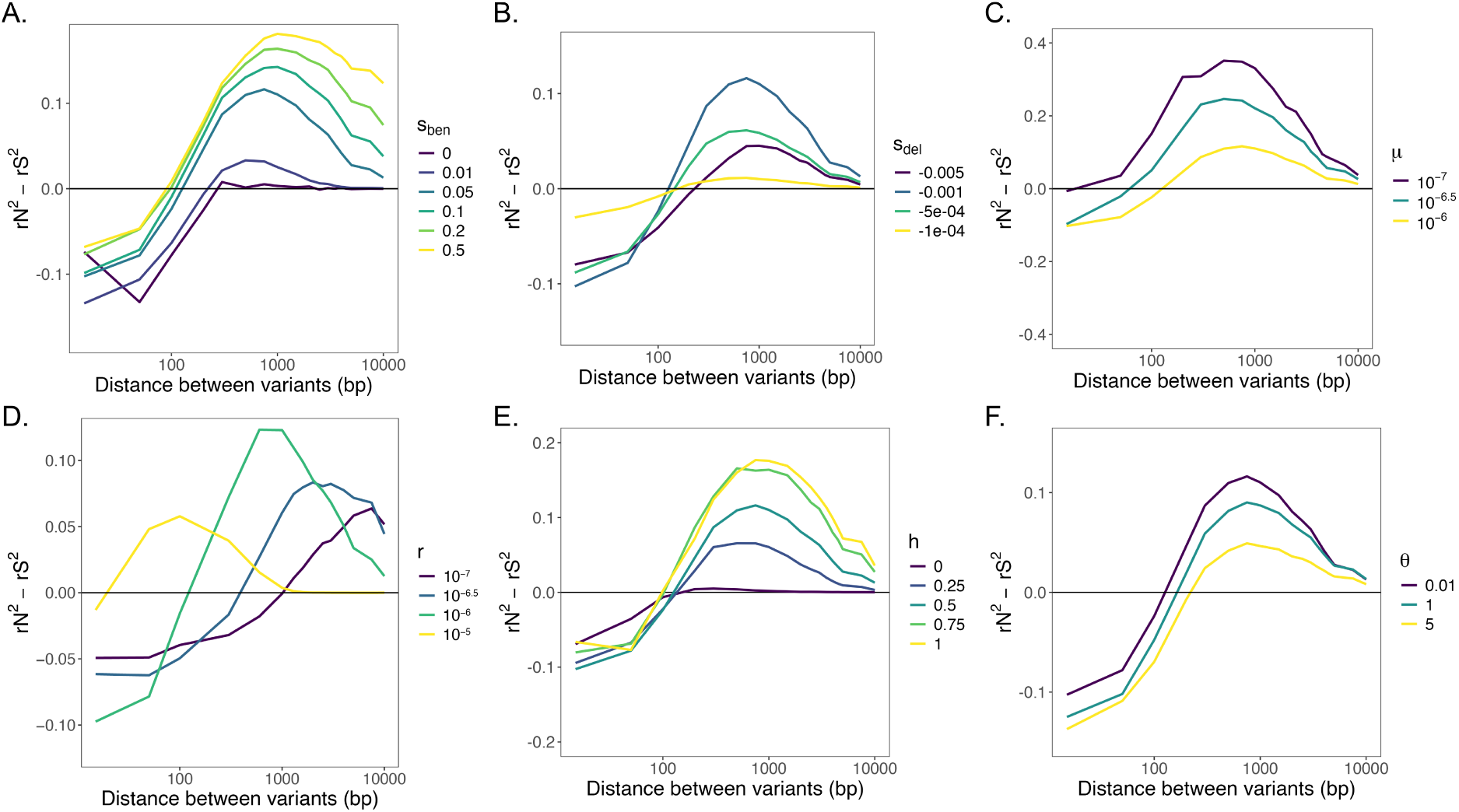
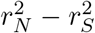 across a range of parameters. Unless otherwise specified, simulations were performed with *Ne* = 10^4^, *s*_*ben*_ = 0.05, *s*_*del*_ = *−*0.001, *µ* = 10^*−*6^, *r* = 10^*−*6^, *h*_*ben*_ = *h*_*del*_ = 0.5, and *θ*_*ben*_ = 0.01. For each combination of parameters, we simulated 1,000 replicates. A. As *s*_*ben*_ increases, 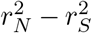 increases. B. As the magnitude of *s*_*del*_ increases, 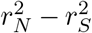 non-monotonically varies at short and farther distances. C. As the mutation rate *µ* increases, 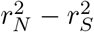 decreases. D. As *r* increases, criss-crossing occurs at shorter distances. E. As the dominance coefficient (*h*_*ben*_ = *h*_*del*_) increases, 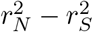 decreases at short distances and increases at farther distances. F. As the adaptive mutation rate, *θ*_*ben*_, increases and the sweeps become soft (*θ*_*ben*_ *>*= 1), 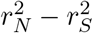 decreases.

Third, the background mutation rate (*µ*_*del*_ = *µ*_*syn*_) impacts the number of deleterious variants circulating in the population, and thus the strength of repulsion (fig. 2C, fig. S4). When the mutation rate is low (*µ* = 10^*−*7^), the probability that two nearby deleterious mutations will arise on the same haplotype is low, and thus 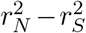 at short distances due to repulsion decreases, resulting in no criss-crossing. As mutation rate increases, the number of nonsynonymous variants increases, thereby increasing the probability nonsynonymous variants will be in repulsion prior to the onset of the sweep, and the entire 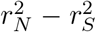 curve shifts downwards, and criss-crossing arises.

Fourth, the recombination rate (*r*) impacts the probability that deleterious variants are broken apart as well as the probability that the beneficial variant is recombined away from its hitchhikers, i.e. the distances at which repulsion and hitchhiking operate (fig. 2D, fig. S4). When recombination is frequent (*r* = 10^*−*5^), criss-crossing occurs at very short distances of 20 bp, which may not be observable in real data if data is not binned finely enough (fig. 2D, fig. S4). As recombination becomes less frequent (*r* = 10^*−*7^), criss-crossing occurs as far as 1000 bps. Additionally, the maximum height of the 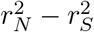 curve occurs at shorter distances, as the distance at which deleterious hitchhikers will be tightly linked to the beneficial variant decreases.

Fifth, the dominance coefficient for nonsynonymous variants (*h*_*ben*_ = *h*_*del*_) impacts the efficiency of selection, and thus the impacts on LD of both repulsion and hitchhiking (fig. 2E). When all nonsynonymous variants are fully recessive (*h* = 0), criss-crossing does not manifest because 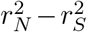 never rises appreciably above zero at any distance. Rather, it remains near zero at most distances and dips below only at short distances (*<* 100 bp). In this fully recessive scenario, selection can act only on variants that are common enough to be inherited in a homozygous state, so low-frequency nonsynonymous variants are effectively neutral and hitchhiking generates no excess LD. Conversely, criss-crossing manifests for any *h >* 0 as selection is felt more readily, and the magnitude increase in 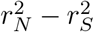 at intermediate distances grows with *h*.

Finally, criss-crossing is a feature of both hard and soft sweeps, simulated with beneficial mutation rates of *θ*_*ben*_ = 0.01 and *θ*_*ben*_ *>* 1, respectively (fig. 2F). However, the magnitude increase in 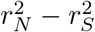 at intermediate distances decreases as *θ*_*ben*_ increases. This is likely because as the number of origins for the sweeping beneficial variant increases, overall LD is attenuated, ^22^ since hitchhikers will be largely private to only one of the sweeping haplotypes.

In sum criss-crossing is a common feature of selective sweeps, with two exceptions when repulsion of deleterious variants cannot be easily felt: low mutation rate (fig. 2C, fig. S4) and complete recessivity of deleterious mutations (fig. 2E).

### 3.3 Repulsion between close-by deleterious variants is masked by hitchhiking near the sweeping locus

Above we found that criss-crossing of 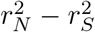 during a sweep may be driven by a mixture of repulsion and hitchhiking. Repulsion should be strongest at short distances between deleterious variants while hitchhiking is strongest for short distances from the sweeping beneficial variant. To disentangle the relative contributions of repulsion vs. hitchhiking, and the distances at which each dominates, we next asked whether the sign of 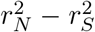 depends on how far apart two deleterious variants are, conditional on their distance from the beneficial variant. We thus decompose the mean 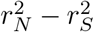 line into pairs that are near the beneficial variant and pairs that are far away from the beneficial variant (fig. S8). We focus the calculation of 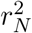 only on pairs of deleterious alleles, omitting pairs that included the beneficial variant, since we are interested in characterizing repulsion between deleterious variants.

When deleterious variants are close to each other (*<* 100 bp apart) and very close to the sweeping beneficial variant (*<* 15 bp away), the effect of hitchhiking is stronger than repulsion leading to 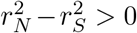 (fig. 3A). As the distance to the beneficial variant increases (*>* 15 bp), we no longer see 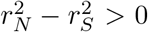 for nearby deleterious variants (fig. S8). As the distance between deleterious variants increases, 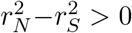, even for pairs not immediately adjacent to the sweeping beneficial variant (fig. 3A). This occurs because as the distance between deleterious variants increases, the probability that recombination can break apart these variants increases, thus decreasing the strength of repulsion. As a consequence, hitchhiking has a stronger effect than repulsion at intermediate distances between deleterious variants, regardless of the proximity to the beneficial variant.

**Figure 3.**
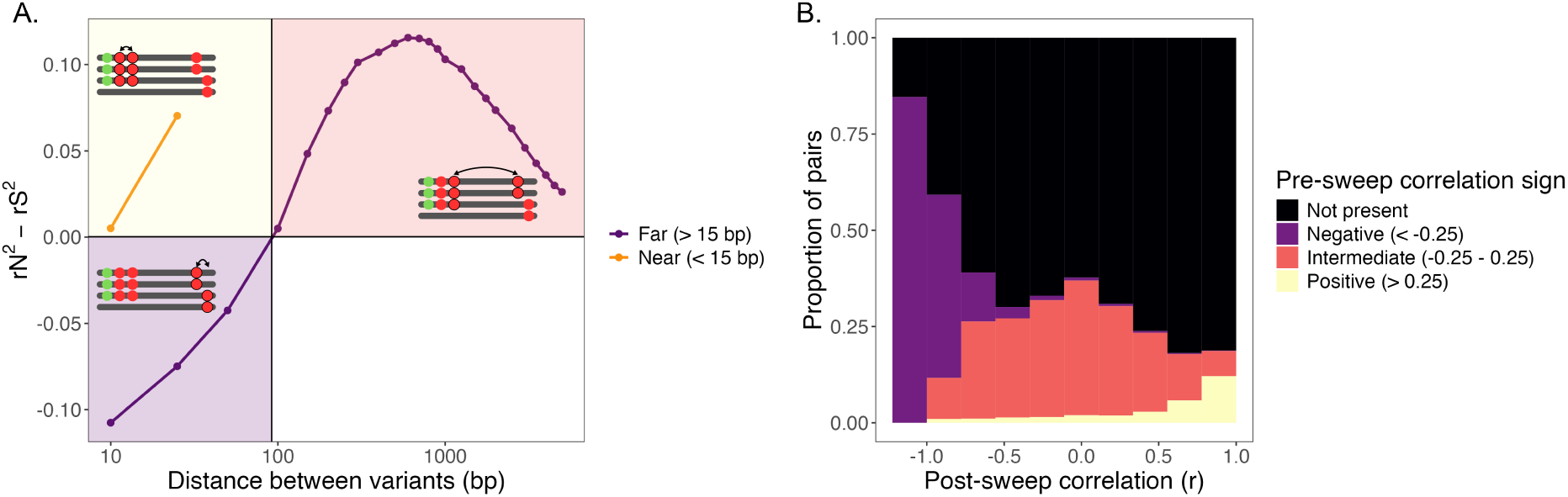
Effects of hitchhiking and repulsion on 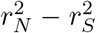 as a function of distance from the beneficial locus. A. Mean 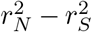, omitting pairs that included the beneficial variant, between intermediate frequency variants (*MAF >* 0.2) following a selective sweep (*s*_*ben*_ = 0.05, *s*_*del*_ = *−*10^*−*3^, *Ne* = 10^4^, *r* = 10^*−*6^, *µ* = 10^*−*6^, *θ*_*ben*_ = 0.01) across 1,000 replicate simulations. Pairs were subdivided based on proximity to the beneficial locus: far pairs are those in which both variants were at least 15 bp away, while near pairs are those in which both variants were at most 15 bp away. Only pairs where both variants were on the same side of the sweeping locus were included. B. Signed correlation coefficients (*r*) were calculated before the onset of positive selection and at the conclusion of the sweep. Pairs in which one or both of the deleterious variants were not present pre-sweep, and thus pre-sweep correlation was impossible to calculate, were labeled as as “Not present.”

Next, to determine whether pairs of alleles in repulsion after the sweep were already in repulsion before the onset of positive selection, we computed the signed correlation coefficient (*r*) between pairs of deleterious variants before vs. after the selective sweep (fig. 3B). Thus far we have used *r*^2^ to quantify correlations. By not squaring the correlation coefficient, we are able to maintain sign and investigate whether intermediate frequency pairs are in repulsion (*r <* 0) or are positively correlated *r >* 0) (Methods). When *r* = *−*1, pairs are perfectly anti-correlated, and every individual in the population has one of the two variants but not both. When *r* = 1, the pairs are perfectly correlated, and every haplotype that bears one of the two variants also has the other. We condition our analysis to only use variants that have reached intermediate frequency post-sweep (*MAF >* 0.2). However, these same variants may be absent prior to the onset of the sweep, in which case it is not useful to calculate pre-sweep correlation. If one or both of the variants were absent prior to the onset of the sweep, the pre-sweep correlation coefficient was labeled “not present” (black). We find that virtually all pairs that were in repulsion pre-sweep (*r*_*pre*_ *< −*0.25) were also negatively correlated post-sweep (*r*_*post*_ *< −*0.5) (fig. 3B). While pairs in strong repulsion (i.e. *r* = *−*1) would generate high *r*^2^ values, this degree of repulsion is rare (fig. S9). On the other hand, the majority of pairs that were tightly positively correlated post-sweep (*r*_*post*_ *>* 0.5) were either already positively correlated pre-sweep (*r*_*pre*_ *>* 0.25) or arose during the sweep (“not present”) (fig. 3B). This demonstrates not only that pairs are already in repulsion prior to the onset of the sweep, but also that hitchhiking is not sufficient to disrupt existing strong repulsion.

### 3.4 Not all sweeps in nature exhibit criss-crossing

Criss-crossing of 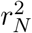 and 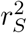 occurs across a range of parameter space in simulations, but the question remains: to what extent do sweeps in nature exhibit this signal? To test this, we analyzed selective sweeps in a dataset of 134 isolates of *Clostridium difficile* for evidence of criss-crossing.

To detect sweeps in *Clostridium difficile*, we used the integrated Linkage Disequilibrium Score (iLDS) Wolff and Garud ^4^, which relies on signals of 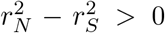. In Figure 4A, iLDS successfully recapitulates known positive controls, including known selective sweeps at *tcdB*, ^23^ and the S-layer cassette. ^15^ Two of the other sweeps identified are known virulence factors, the *fli* operon ^16^ and the PilZ domain, ^24^ while the two remaining sweeps are both in ABC type 3 transporters, which have been implicated in drug resistance.^25^

We note that the iLDS statistic is always positive, as the difference between 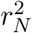 and 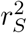 is squared in the statistic. ^4^ This has the potential to obfuscate signatures of criss-crossing, which could generate negative values of 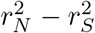. To investigate if specific sweeps exhibit criss-crossing, it is thus necessary to plot 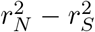 for each sweep. We calculated 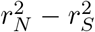 for all intermediate frequency (*MAF ≥* 0.2) variants within a distance of 10^4.5^ bps from the center of the sweep (Methods).

We find that some, but not all, sweeps exhibit criss-crossing of LD (fig. 4B). The sweeps in the first ABC type 3 transporter, *tcdB*, and the PilZ domain unambiguously demonstrate criss-crossing of LD, consistent with our simulations. On the other hand, the sweeps in the *fli* gene and the S-layer cassette exhibit 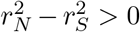 across all distances, despite binning the data finely at short distances (Methods).

We next asked what process could generate an absence of criss-crossing, or, 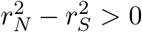 across all distances. In our simulations, only low mutation rate produced an elevation in 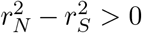. However, the scaled mutation rate in this scenario was 4 *\** 10^*−*3^ (*θ* = 4*N*_*e*_*µ* = 4 *\** 10^4^ *\** 10^*−*7^). By contrast in *C. difficile*, we find that the average nucleotide diversity in neutral regions (Methods) is *π ≈* 1.6 *\** 10^*−*2^, which is in the regime of the mutation rate that does generate criss-crossing in simulations (fig. 2C).

We found also for high recombination rates (*ρ/θ >* 1) criss-crossing occurs only at very short distances (*≈* 20 bp). Since the smallest distance bin we used (30 bp) was greater than this distance, it is possible criss-crossing is unobservable in our data due to insufficient SNP density. However, the recombination rates that produced criss-crossing at short distances (fig. 4), is higher than that observed in the data: Didelot and Wilson ^26^ estimated *ρ/θ* = 0.3 for *C. difficile*.

**Figure 4.**
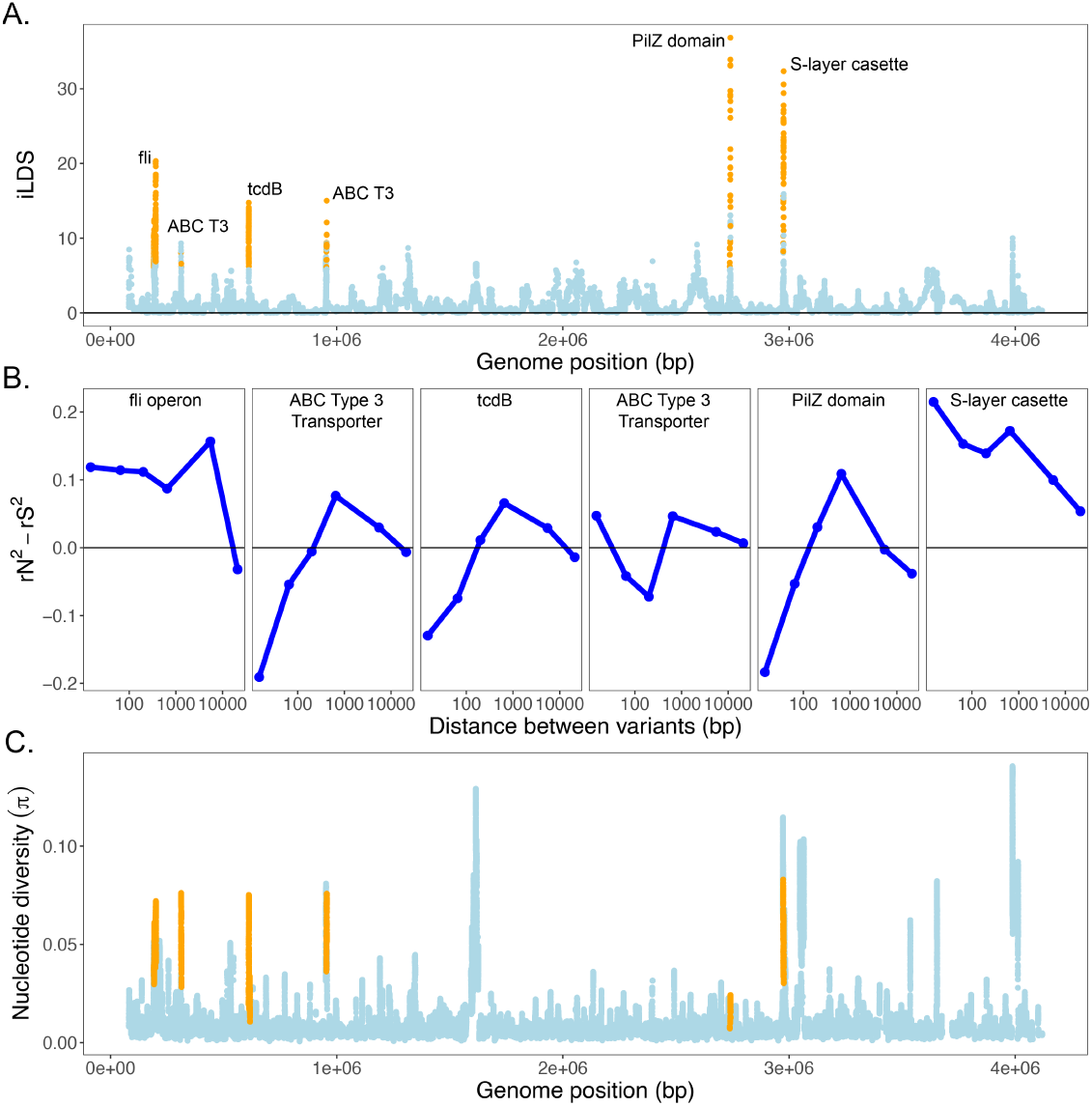
Only some sweeps in nature exhibit criss-crossing. A. iLDS scan of 134 isolates of *Clostridium difficile*, using an analysis window roughly corresponding to 10^4.5^ bps (Methods), recapitulating the scan from. ^4^ Each point corresponds to a genomic window centered around a single intermediate frequency nonsynonymous SNP. Windows harboring a statistically significant selective sweep by iLDS are colored orange, whereas non-significant windows are colored blue. B. Mean 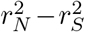 for each of the six significant sweeps identified by iLDS. *r*^2^ was calculated for all intermediate frequency variants (*MAF >* 0.2) within 10^4.5^ bp of the central locus of the sweep. 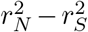 was binned at distances of 30, 100, 500, 1000, 10000, 32000 such that there were at least 5 pairs per bin. C. Mean average nucleotide diversity (*π*), calculated in sliding windows of 1000 bp long. Windows overlapping sweep regions identified by iLDS are colored orange.

One possibility that could result in lack of criss-crossing is the adaptive architecture of the sweeps. Previous work has shown that complex, multi-locus adaptations and introgression can generate elevated 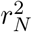 relative to 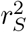 Stolyarova et al. ^27^, Arnold et al. ^28^. Epistasis can cause nearby deleterious variants to become tightly linked, leading to elevated 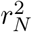 compared to 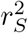. ^27^ Additionally, adaptive introgression, in which adaptive material is acquired via horizontal gene transfer from a distant lineage, causes a high density of variants at short distances to be simultaneously acquired, ^28^ also leading to elevated 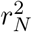 compared to 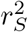.

Both of the sweeps that do not exhibit criss-crossing in *C. difficile* have been previously shown to be complex, multi-locus adaptations. ^15,16^ However, the sweep at *tcdB* is also complex, yet still exhibits criss-crossing. The adaptation at *tcdB* is a multi-locus adaptation made up of four functional domains that recombine independently.^23^ When we account for domain boundaries at *tcdB*, we see 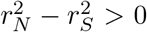 across all distances for pairs within domain boundaries, while we still see criss-crossing for pairs in different domains (fig. S10).

Moreover, we find that in *C. difficile*, the majority of sweeps identified by iLDS, including the two that do not exhibit criss-crossing, show elevated *π* relative to the genome-wide mean (fig. 4C). Hard sweeps of single locus adaptations, by contrast are expected to decrease *π* in the vicinity of the sweep, ^29^ while both introgression and complex adaptive architecture may increase *π* Laurin and Garud ^30^.

The patterns observed at these sweeps exhibiting lack of criss crossing are consistent with Stolyarova et al. ^27^ and Arnold et al. ^28^. However, neither Stolyarova et al. ^27^ or Arnold et al. ^28^ considered the effect of deleterious hitchhikers in conjunction with these other processes. Thus, in the next section, we test the hypothesis that complex adaptive architecture and adaptive introgression may reduce the effect of repulsion on patterns of LD and instead result in 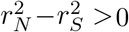 across all distances.

### 3.5 Introgression and epistasis can result in a lack of criss-crossing

Here, we test the hypothesis that introgression of adaptations or complex, epistatic adaptations can result in a lack of criss-crossing even when there are hitchhiking deleterious variants. To do so, we simulated a selective sweep where the adaptation lies on an adaptive fragment of *n*_*loci*_ nonsynonymous variants that is beneficial when all *n*_*loci*_ variants are found on the same haplotype (*s*_*together*_ = 0.1) (Methods). As was the case before, we allow for neutral and deleterious (*s*_*del*_ = *−*10^*−*3^) hitchhikers to evolve with rate *µ* = 10^*−*6^ and a recombination rate of *r* = 10^*−*6^.

First, to compare the epistasis and adaptive introgression scenarios with that of a single locus sweep, we plot *n*_*loci*_ = 1 in Figure 5. Replicating results from above, 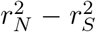 exhibits criss-crossing (fig. 5, black line).

**Figure 5.**
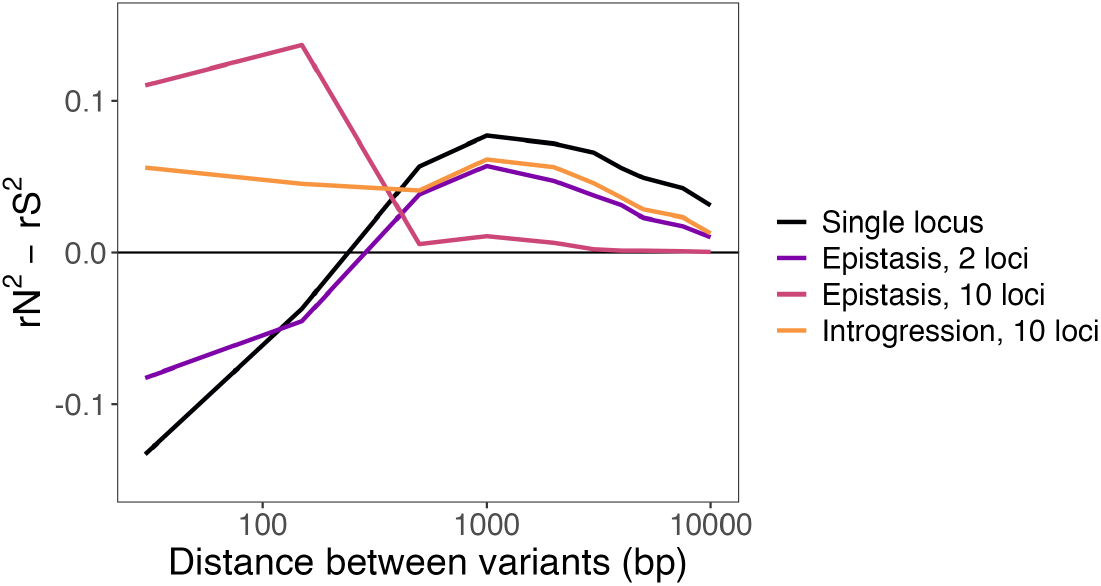
Effects of adaptative introgression and complex adaptation on 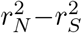. When *n*_*loci*_ = 1, this mirrors evolution via de novo mutation, and criss-crossing is observed (black line). To simulate epistasis, we consider models where *n*_*loci*_ *>* 1 nonsynonymous variants interact such that they are each deleterious alone (*s*_*alone*_ = *−*0.001) and only become adaptive when all mutations are present on the same haplotype. When there are two epistatically interacting loci, criss-crossing is observed (purple), while when there are *n*_*loci*_ = 10 criss-crossing is not observed (pink). In the introgression model (orange), the center-most mutation is beneficial while the remaining *n*_*loci*_ *−* 1 mutations are deleterious. Once again, criss-crossing is not observed.

Next, to simulate epistasis, we increase *n*_*loci*_ *>* 1. In this scenario, each mutation is deleterious alone (*s*_*alone*_ = *s*_*del*_ = *−*0.001) and only becomes beneficial when all *n*_*loci*_ adaptations are present on the same haplotype. We find that when there are only two epistatically interacting loci, 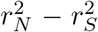 still exhibits criss-crossing (fig. 5, purple line). However, when there are 10 epistatically interacting loci, 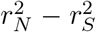 is elevated above zero across all distances (fig. 5, pink line).

Finally, to simulate adaptive introgression, we consider an adaptive fragment with *n*_*loci*_ = 10 mutations where only the center-most mutation is beneficial while the remaining *n*_*loci*_ *–* 1 mutations are deleterious hitchhikers. In this case, we again find that 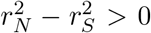 across all distances (fig. 5, orange line).

Thus, models of both epistasis with large numbers of interacting loci and adaptive introgression can mask repulsion between deleterious hitchhikers and lead to a lack of criss-crossing in 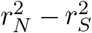.

## 4 Discussion

During a selective sweep, deleterious nonsynonymous variants can hitchhike to high frequency. ^4–6^ Here, we show that hitchhiking and repulsion have opposing effects on LD at a sweep, leading to elevation in LD between pairs of intermediate frequency nonsynonymous vs synonymous variants at some, but not all, distances (fig. 1C). More specifically, we find that on average hitchhiking generates elevated nonsynonymous LD 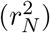 relative to synonymous LD 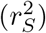 at intermediate distances between variants, but repulsion causes 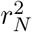 to be less than 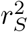 at short distances. The relative strength of hitchhiking vs. repulsion is not just a function of distance between variants but also distance to the beneficial variant: hitchhiking can have a larger effect on LD than repulsion of close-by deleterious pairs only when they are very close to the sweeping beneficial variant. Together, these dynamics create a characteristic criss-crossing of LD as a function of distance between variants.

The dynamics of hitchhiking deleterious variants have been an area of focus in the field: others have established that deleterious hitchhiking can inhibit selective sweeps ^31^ and increase the prevalence of intermediate frequency deleterious variants. ^6^ However, the simultaneous effect of hitchhiking and repulsion between deleterious variants has not yet been explored. Our work recapitulates the findings of Wolff and Garud ^4^, demonstrating that hitchhiking alone is sufficient to explain on average 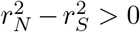. By using finer binning with respect to distance (fig. S6), we discover that this elevation is non-uniform, as repulsion generates 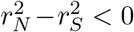 for short distances between variants. Wolff and Garud ^4^ leveraged this signal of elevated nonsynonymous LD to identify selective sweeps with the statistic the integrated Linkage Disequilibrium Score (iLDS). Our finding of criss-crossing may improve the resolution of iLDS, as the statistic was calibrated to detect overall 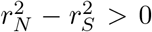. Since criss-crossing of LD will result in 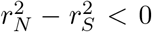 at short distances, the extent of the increase of 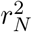 over 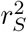 in a sweep may be muted, resulting in some sweeps potentially being missed by iLDS. In the future, accounting for signatures of repulsion in iLDS and other LD based selection scan statistics may improve their sensitivity.

While criss-crossing occurs across a wide range of biologically relevant parameter space (fig. 2, S2, S3, S4), we find that it does not uniformly manifest in data. In particular, two sweeps in *C. difficile* that do not exhibit criss-crossing have been previously shown to be complex, multi-locus adaptations. ^15,16^ Moreover, most of the sweeps identified in *C. difficile* coincide with regions of elevated nucleotide diversity, suggestive of adaptive introgression. ^30^ Consistent with this, our simulations and previous work ^27,28^ show that introgression and complex, multi-locus sweeps (i.e. epistasis) can generate 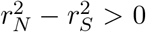 across all distances. We note that a combination of forces likely generates 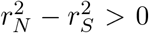 in real data: hitchhiking of deleterious alleles alongside complex adaptations and, possibly, introgression. Understanding the specific genetic architectures of sweeps in real data will require future work.

The presence of criss-crossing does not necessarily imply a lack of complex adaptation or introgression. For example, independently recombining domains in *tcdB* generated criss-crossing for the average 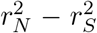 across the whole gene, despite *tcdB* being a complex adaptation. ^23^ Additionally, we find that only two epistatically interacting loci were not sufficient to mask signatures of repulsion (fig. 5). Thus, it may be that only specific architectures of complex adaptation are able to result in 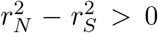 across all distances. As such, the presence of criss-crossing does not preclude that the adaptation is complex, especially in cases of domain mosaicism, few interacting loci, or existing deleterious variation already in repulsion.

In summary, our findings characterize the interaction between two population genetic forces: hitchhiking and repulsion. In the future, research that leverages criss-crossing, or a lack thereof, to characterize the adaptive architecture of selective sweeps in nature could help us better understand the genetic architecture of sweeps across diverse systems.

## Supporting information

All Supplemental Figures

## 5 Data availability

All code and data is available on GitHub (https://github.com/garudlab/hitchhiking_repulsion_ld).

Haplotype data for C. difficile were obtained from UHGG v1^18^ available via.

## 6 Acknowledgments

## 7 Funding

This work was funded by the US National Institutes of Health National Institute of General Medical Sciences awards R35GM151023 to NRG and R35GM119856 to KEL, a National Science

Foundation CAREER award (no. 2240098) to NRG, and the UCLA QCBio Collaboratory Award to MW. We would like to thank all members of the Garud Laboratory, especially Richard Wolff, Peter Laurin, and Michael Wasney, for their feedback.

## 8 Conflicts of interest

The authors have no conflicts of interest to report.

## Notes

https://github.com/garudlab/hitchhiking_repulsion_ld

### Competing Interest Statement

The authors have declared no competing interest.

