## Supplementary material for "The effects of hitchhiking and repulsion of deleterious alleles on linkage disequilibrium during a selective sweep": All Supplemental Figures

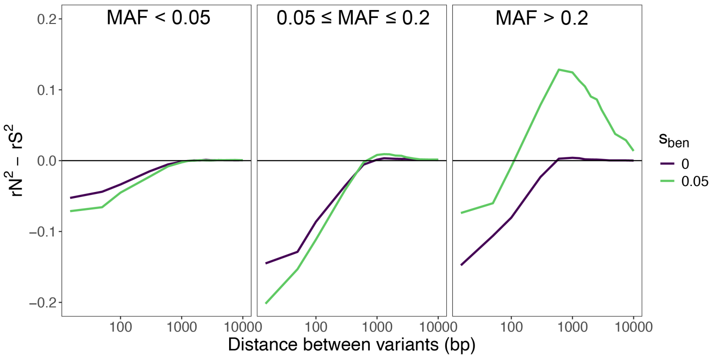


Supplemental Figure 1: Effect of minor allele frequency (MAF) on criss-crossing. Mean $r_N^2 - r_S^2$ curves constructed from at least 1,000 replicate simulations. Each simulation was conducted with $Ne = 10^4$, $r = 10^{-6}$, $\mu = 10^{-6}$, and $\theta_{ben} = 0.01$, and $s_{del} = -10^{-3}$.

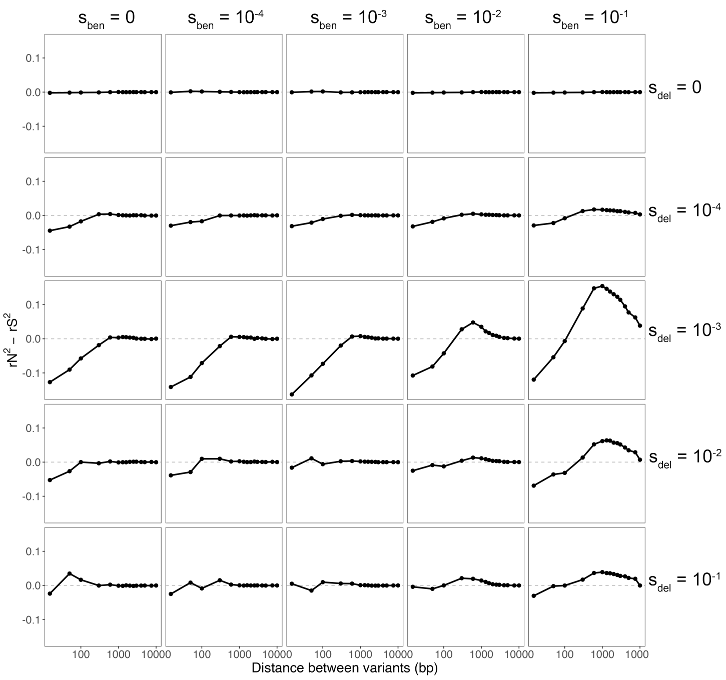


Supplemental Figure 2: Criss-crossing across strength of beneficial selection ($s_{ben}$) and strength of deleterious selection ($s_{del}$). Mean $r_N^2 - r_S^2$ curves constructed from at least 1,000 replicate simulations such that each point plotted represents a mean of at least 100 pairs. Each simulation was conducted with $Ne = 10^4$, $r = 10^{-6}$, $\mu = 10^{-6}$, and $\theta_{ben} = 0.01$. Criss-crossing occurs when $s_{ben} > |s_{del}| > 1/Ne$. When $s_{del} = 0$, there is no difference between synonymous and nonsynonymous variants, and $r_N^2 - r_S^2 = 0$ across all distances for all values of $s_{ben}$. When $1/Ne < |s_{del}| < 10^{-1}$, repulsion generates $r_N^2 - r_S^2 < 0$ at short distances. When the strength of deleterious selection is too strong ($|s_{del}| = 10^{-1}$), deleterious hitchhikers are unable to drift to intermediate frequency and we no longer see $r_N^2 - r_S^2 < 0$ due to repulsion.

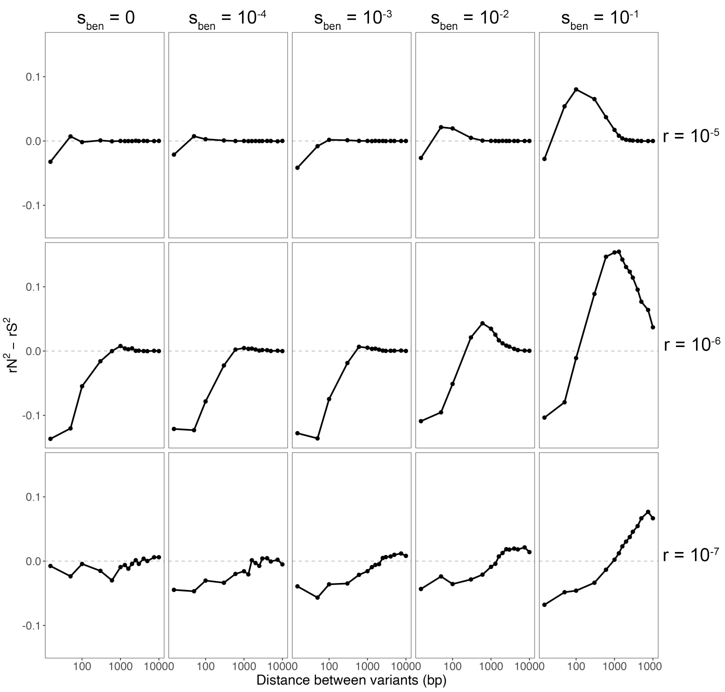


Supplemental Figure 3: Criss-crossing across strength of beneficial selection ($s_{ben}$) and recombination rate ($r$). Mean $r_N^2 - r_S^2$ curves constructed from at least 1,000 replicate simulations, such that each point plotted represents a mean of at least 100 pairs, where $Ne = 10^4$, $s_{del} = -10^{-3}$, $\mu = 10^{-6}$, and $\theta_{ben} = 0.01$. When $s_{ben} \leq s_{del}$, while there is a decrease in $r_N^2 - r_S^2 < 0$ at short distances due to repulsion, there $r_N^2 - r_S^2$ never exceeds 0. When recombination is frequent ($r = 10^{-5}$), $r_N^2 - r_S^2 < 0$ only at very short distances $< 15$ bp apart for all cases where $s_{ben} > s_{del}$. When $r = 10^{-6}$, $r_N^2 - r_S^2 < 0$ for distances $< 200$ bp apart. When $r = 10^{-7}$ and recombination is very infrequent, $r_N^2 - r_S^2 < 0$ for distances $< 750$ bp apart.

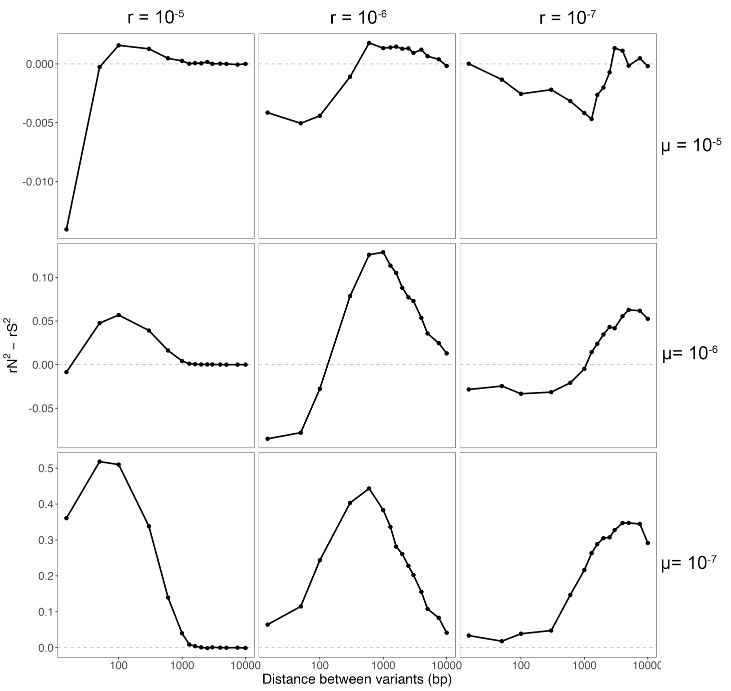


Supplemental Figure 4: Criss-crossing across recombination rate ($r$) and background mutation rate ($\mu$ ). Mean $r_N^2 - r_S^2$ curves constructed from at least 1,000 replicate simulations, such that each point plotted represents a mean of at least 100 pairs, where $Ne = 10^4$, $s_{del} = -10^{-3}$, $s_{ben} = 0.05$, and $\theta_{ben} = 0.01$. Criss-crossing occurs for intermediate values of mutation and recombination rate. When the mutation rate is high ($\mu = 10^{-5}$), deleterious nonsynonymous variants are more readily able to drift to intermediate frequency, and there is little difference between $r_N^2$ and $r_S^2$. Conversely, when the mutation rate is low ($\mu = 10^{-7}$), few deleterious nonsynonymous mutations are able to drift to intermediate frequencies, resulting in no $r_N^2 - r_S^2 < 0$ at short distances due to repulsion.

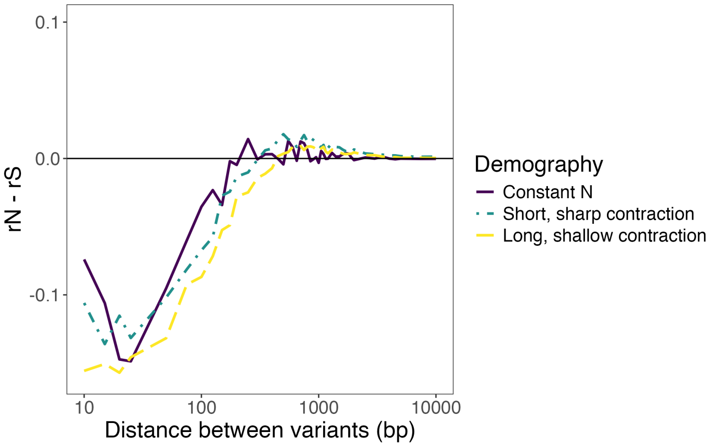


Supplemental Figure 5: $r_N^2 - r_S^2$ under demographic contractions. In the "constant Ne" model (purple), the population size was held constant at $N_e = 10^4$ and simulations ended after $10N_e$ generations. Under both contractions, an instantaneous population size change occurred occurred following the burn-in. In the "short, sharp contraction" model, the population size was reduced to $1/10 * N_e$ and simulations continued for an additional 20 generations. In the "long, shallow contraction" model, the population size was halved and the simulations continued for an additional 5000 generations. Under all three demographic scenarios, we do not see criss-crossing: $r_N^2 < r_S^2$ at short distances, and decays to become equivalent at farther distances ($r_N^2 - r_S^2 \approx 0$). LD was calculated for intermediate frequency ($MAF > 0.2$) variants.

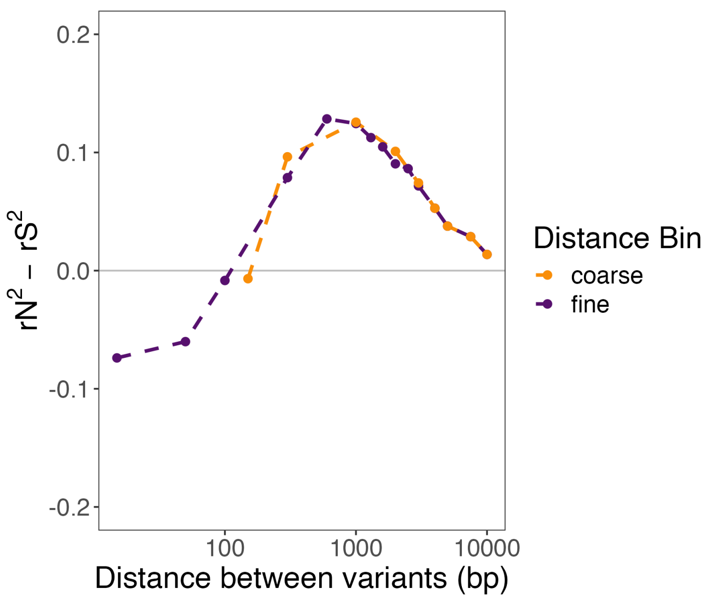


Supplemental Figure 6: Effect of coarse vs fine binning on visualizing criss-crossing. Criss-crossing of LD is only apparent when distance between variants is binned finely at short distances. Using the same underlying simulations, we calculated mean $r^2_N - r^2_S$ across 1,000 replicate simulations where $s_{ben} = 0.1$, $s_{del} = -10^{-3}$, an $Ne = 10^4$. When distance is binned finely at short distances (i.e. breaks at 15, 50, 100, and 300), $r^2_N - r^2_S < 0$ at short distances between variants (purple). When distance is binned coarsely (i.e. the smallest bin includes pairs $< 150$ bp apart), as was the case in Wolff and Garud [4], it appears that $r^2_N - r^2_S \geq 0$ across all distances, and we no longer see criss-crossing.

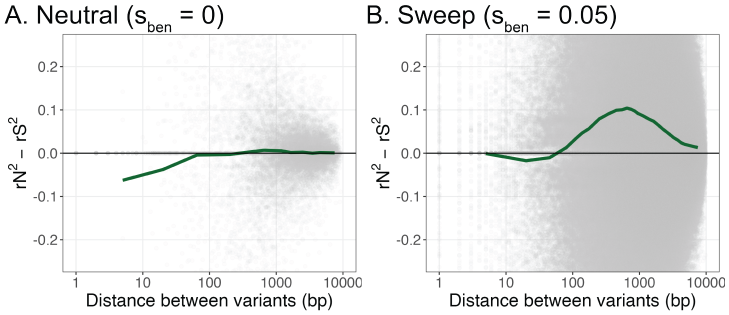


Supplemental Figure 7: Effect of allele frequency matching on criss-crossing. $r_N^2 - r_S^2$ calculated for synonymous and nonsynonymous pairs that were precisely matched in terms of both allele frequency and distance. Each grey point corresponds to a nonsynonymous pair and all equivalent pairs of synonymous variants with precisely the same a) distance apart, and b) allele frequencies (i.e. $f_{N1} = f_S 1 \& f_{N2} = f_{S2}$). If no synonymous pairs matched all criteria, the nonsynonymous pair was discarded. Finally, the moving average of the frequency-distance matched pairs was calculated (green line). A. When $s_{ben} = 0$, and there is no hitchhiking, we find that the moving average $r_N^2 - r_S^2$ does not exhibit criss-crossing, as the mean $r_N^2 - r_S^2$ is not greater than 0 at farther distances. B. When $s_b en = 0.05$, we find that the moving average line still exhibits criss-crossing, similar to the binned data.

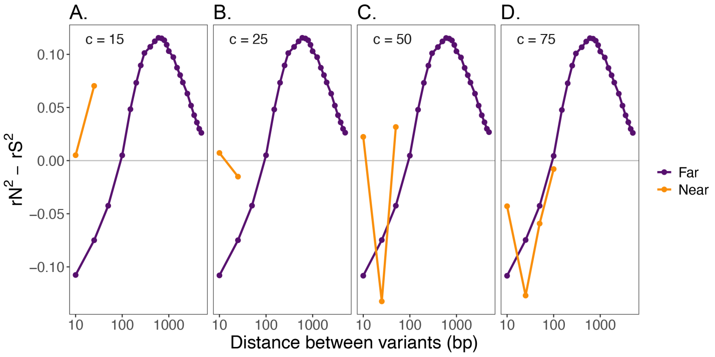


Supplemental Figure 8: Effect of distance of deleterious pairs from the beneficial variant on $r_N^2 - r_S^2$. The x axis represents distance between deleterious variants. Here $r_N^2 - r_S^2$ was computed omitting pairs that included the beneficial variant, between intermediate frequency variants ($MAF > 0.2$) following a selective sweep ($s_{ben} = 0.05$, $s_{del} = -10^{-3}$, $Ne = 10^4$). Only pairs where both variants were on the same side of the sweeping locus were included. Pairs where both deleterious variants were $> c$ bp away from the beneficial variant were labeled as "Far" (purple), while pairs that were less than $c$ bp away from the beneficial variant were labeled "Near" (orange). If both variants in the pair did not belong to the same group (i.e. one was "Far" and the other was "Near"), the pair was omitted. A-D. The value of the $c$, or the distance to beneficial variant cut-off, was varied.

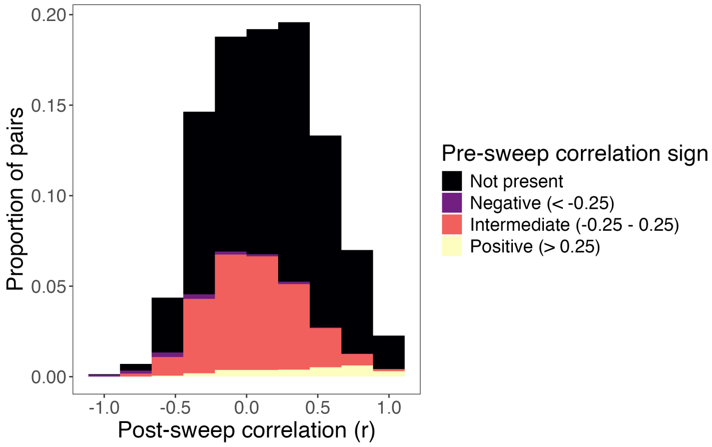


Supplemental Figure 9: Distribution of signed correlation coefficients $(r)$ between pairs of deleterious nonsynonymous intermediate frequency variants $(MAF > 0.2)$. Here $(s_{ben} = 0.05$, $s_{del} = -10^{-3}$, $Ne = 10^4$, $r = 10^{-6}$, $\mu = 10^{-6}$, $\theta_{ben} = 0.01)$ across 1,000 replicate simulations. Signed correlation coefficients $(r)$ were calculated before the onset of positive selection and at the conclusion of the sweep. Pairs in which one or both of the deleterious variants were not present pre-sweep, and thus pre-sweep correlation was impossible to calculate, were labeled as as “Not present.”

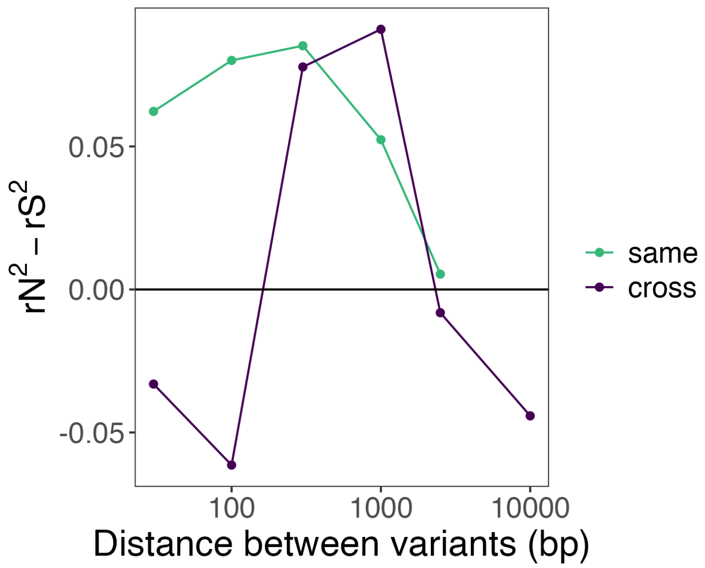


Supplemental Figure 10: Mean $r_N^2 - r_S^2$ for intermediate frequency ($MAF >= 0.2$) variants in the *tcdB* gene. The *tcdB* gene is composed of four different domains.[23] Pairs were thus subdivided based on whether both variants were in the same domain (green) or variants were across domains (dark purple). Within domains, we see $r_N^2 - r_S^2 > 0$ even at short distances, consistent with our simulations of adaption via complex, multi-locus architecture. Pairs found across domains exhibit $r_N^2 - r_S^2 < 0$ at short distances.
